# Effects of repeated developmental GLP-1R agonist exposure on adult behavior and hippocampal structure in mice

**DOI:** 10.1101/2023.04.21.537724

**Authors:** Catherine Cerroni, Alex Steiner, Leann Seanez, Sam Kwon, Alan S. Lewis

**Author notes:** **Corresponding author:** Alan S. Lewis, MD, PhD, Assistant Professor of Psychiatry and Behavioral Sciences, Vanderbilt University Medical Center, 465 21st Ave. S., MRBIII Room 6140B, Nashville, TN 37240. Denotes co-first authors (Catherine Cerroni, Alex Steiner).

## Abstract

Glucagon-like peptide-1 receptor (GLP-1R) agonists are common type 2 diabetes medications that have been repurposed for adult chronic weight management. Clinical trials suggest this class may also be beneficial for obesity in pediatric populations. Since several GLP-1R agonists cross the blood-brain barrier, it is important to understand how postnatal developmental exposure to GLP-1R agonists might affect brain structure and function in adulthood. Toward that end, we systemically treated male and female C57BL/6 mice with the GLP-1R agonist exendin-4 (0.5 mg/kg, twice daily) or saline from postnatal day 14 to 21, then allowed uninterrupted development to adulthood. Beginning at 7 weeks of age, we performed open field and marble burying tests to assess motor behavior and the spontaneous location recognition (SLR) task to assess hippocampal-dependent pattern separation and memory. Mice were sacrificed, and we counted ventral hippocampal mossy cells, as we have recently shown that most murine hippocampal neuronal GLP-1R is expressed in this cell population. We found that GLP-1R agonist treatment did not alter P14-P21 weight gain, but modestly reduced adult open field distance traveled and marble burying. Despite these motor changes, there was no effect on SLR memory performance or time spent investigating objects. Finally, we did not detect any changes in ventral mossy cell number using two different markers. These data suggest developmental exposure to GLP-1R agonists might have specific rather than global effects on behavior later in life and that extensive additional study is necessary to clarify how drug timing and dose affect distinct constellations of behavior in adulthood.

## INTRODUCTION

Agonists of the glucagon-like peptide-1 receptor (GLP-1R) have been in common clinical use to treat diabetes for almost two decades. Their safety and ease of dosing make them largely acceptable to patients and positions them for repurposing to treat additional indications. Along these lines, two GLP-1R agonists, semaglutide and liraglutide, have shown clinically meaningful effects on weight loss in adults with overweight or obesity [1-4], resulting in FDA approval for this indication. Clinical trials have also shown benefit for the GLP-1R agonists liraglutide [5], semaglutide [6], and exenatide [7] in treating pediatric obesity. As obesity in children is prevalent and confers significant morbidity, it is of great public health importance to identify new methods to treat this population.

Several GLP-1R agonists cross the blood-brain barrier [8, 9] where they act on G protein-coupled GLP-1Rs expressed in numerous brain regions. Extensive previous work has shown that GLP-1R agonists as well as endogenous GLP-1 released from the nucleus tractus solitarius influence cognitive, appetitive, and homeostatic processes in adult animals (reviewed in [10]).However, much less is known as to how activation of GLP-1Rs during normal postnatal brain development influence brain structure and function in adulthood. Mice exposed to GLP-1R agonists in utero do not show changes in cognition or locomotor activity as adults [11], and most studies of GLP-1R agonist delivery during rodent postnatal development have focused on their neuroprotective effects in models of neurological disease [12, 13] or in regulating metabolism [14-16]. Our question is important for at least two reasons. First, understanding the role for GLP-1R signaling in brain development is significant for elucidating fundamental neurobiological mechanisms. Second, because of the prevalence of childhood obesity, the number of children potentially eligible for treatment with GLP-1R agonists if they are ultimately proven beneficial for this indication is substantial, and long-term effects on development cannot be identified in standard clinical trials.

In this study, we begin to address the complex question of how GLP-1R agonist treatment during development influences brain structure and function later in life. We treated mice systemically with the GLP-1R agonist exendin-4 (a naturally occurring version of the FDA-approved pharmaceutical exenatide) or saline between postnatal day 14 (P14) and P21. This time period corresponds to a proposed period of *Glp1r* expression increase in mouse brain [11]. Mice were then weaned, allowed to develop uninterrupted, tested in behavioral experiments starting at 7 weeks of age, and then sacrificed for immunohistochemistry. While GLP-1Rs are expressed in many brain areas, we focused our study on the hippocampus, where we have recently shown in mice that GLP-1Rs are highly expressed in mossy cells (MCs) of the dentate gyrus (DG), especially MCs of the ventral DG [17]. Our study’s main findings were modest reductions in locomotor measures, no significant changes in a sensitive test of hippocampal-dependent learning and memory, and no significant change in ventral MC number. These results suggest early developmental GLP-1R activation may have behavior-specific effects that warrant extensive future study to clarify.

## MATERIALS AND METHODS

### Animals

Male and female C57BL/6 mice were bred at Vanderbilt University and group housed under standard conditions with a 12-hour light/dark cycle with ad libitum access to food and water. 22q11.2 deletion syndrome heterozygous mice (C57BL/6-Del(16Dgcr2-Hira)1Tac, Model 11026) [18] and their wildtype littermates on C57BL/6 background were originally purchased from Taconic Biosciences (Germantown, NY), bred at Vanderbilt, genotyped according to standard vendor protocols, and housed in the same conditions as above. All procedures were approved by the Vanderbilt Institutional Animal Care and Use Committee.

### Drug administration

Exendin-4 acetate was purchased from Cayman Chemical Company (catalogue no. 11096, Ann Arbor, MI) and dissolved in saline. From P14 to P21, pups were injected with exendin-4 (0.5 mg/kg) or saline twice daily. Mice were then weaned as usual and allowed to develop uninterrupted until behavioral study.

### Behavioral studies

#### General

Mice were brought from the vivarium to the testing room and allowed to habituate for at least 1 h. All behavioral testing occurred between 9 am and 5 pm. Saline- and exendin-4-treated mice were always tested on the same days, and all tests were videotaped. The order of behavioral tasks was open field, then marble burying, then spontaneous location recognition.

#### Open field test

Performed as previously described [19]. Briefly, animals were placed in a 61 × 61 cm arena and allowed to freely explore for 20 min. Total distance traveled, time spent in the center, and crosses into the center were measured using ANY-maze version 6.1 (Stoelting, Wood Dale, IL).

#### Marble burying test

We placed ten marbles approximately 2-4 cm apart in 5-10 cm of bedding in each animal’s cage [20]. After room acclimation for 1 h, mice were placed in the cages for 30 min. Afterward, the experimenter counted the number of marbles buried for each mouse. Observations were blinded and a marble was counted as buried if two-thirds of the marble was covered by bedding.

#### Spontaneous location recognition (SLR) test

The SLR test was conducted using the similar-SLR (“s-SLR”) protocol outlined by Reichelt et al. [21]. Animals were habituated to a circular arena. During the sample phase, two identical objects were placed 72° apart while a third identical object was placed opposite the other two. Animals explored for 10 min during the sample phase and then were placed back in their home cage. Three h later mice underwent the test phase. The one object originally placed across from the other two remained in the same place (“familiar location”), while another identical object was placed in between the previous locations of the other two identical objects (“novel location”). Animals explored the two identical objects in the test phase for 5 min. All objects and arenas were cleaned before each trial. Sample and test phases were recorded and scored blindly by the experimenter. Results were expressed as a discrimination index (DI) from the test phase, calculated using the formula DI = [time(novel) - time(familiar)] divided by [time(novel) + time(familiar)].

### Immunohistochemistry

Perfusion and tissue slicing were performed exactly as previously described [19]. Free-floating sections were permeabilized and blocked in 0.3% Triton X-100 and 3% normal donkey serum (Jackson ImmunoResearch, West Grove, PA) in PBS for 2 h at room temperature. Sections were incubated in rabbit anti-GluR2/3 (AB1506, Millipore Sigma, Burlington, MA, 1:200) and mouse anti-calretinin (MAB1568, Millipore Sigma, 1:1000) overnight at 4 °C. Sections were washed three times in PBS for 10 min each and then incubated in donkey anti-rabbit Alexa 488 and donkey anti-mouse Alexa 647 (Jackson ImmunoResearch, 1:1000) for 2 h at room temperature. Slices were washed in PBS, incubated with 4′,6-diamidino-2-phenylindol (DAPI, Millipore, 1:5000) for 5 min, washed again in PBS and mounted using Fluoromount-G (Electron Microscopy Sciences, Hatfield, PA).

### Microscopy and image quantification

Images were acquired with an LSM Zeiss 880 confocal microscope (Zeiss, White Plains, NY) and Zeiss Zen software. The total number of calretinin and GluR2/3 positive cells within the DG hilus were counted and averaged across 2 sections per mouse to obtain a single value for each mouse, which were then averaged across groups. All image analysis was performed blind to treatment using Fiji [22].

### Statistical analysis

Unpaired t tests with Welch’s correction were used to compare two groups. Mixed two-way analysis of variance (ANOVA) was used to compare four groups. One-sample t tests were used to compare DI versus 0. Significance was set at p < 0.05. All tests were two-tailed. Analyses and data plots were performed using Prism 9 (GraphPad, San Diego, CA). Standard error of the mean is shown by all error bars. Statistical analyses are performed on male and female combined groups. Male and female data are also shown disaggregated. However, statistical analyses were not performed on disaggregated data as the studies were not powered to formally detect sex differences.

### Data availability

All data from this study are available in **Supplemental File 1**.

## RESULTS

### Experimental timeline and dose selection

The experimental timeline is shown in **Figure 1a**. Beginning at P14, male and female mouse pups were treated with exendin-4 (0.5 mg/kg) i.p. twice daily or saline until P21. We chose this dose based on a previous study in which mouse pups beginning at P7 were repeatedly administered exendin-4 at 0.5 mg/kg to test its effects on hypoxic-ischemic brain injury outcomes [12]. This study reported neuroprotective benefits without effects on blood glucose levels, blood counts and chemistries, organ histopathology, or inflammatory markers. This is a high dose of exendin-4, which we felt was advantageous for our study’s purposes as it might enhance our likelihood of detecting subsequent effects during adulthood. We did not detect an effect of exendin-4 on pup weight gain (**Figure 1b-d**). Beginning at approximately 7 weeks of age, mice underwent two tests reflective of motor behavior (open field and marble burying tests), and one test of hippocampal-dependent pattern separation and memory (SLR) [21]. At the completion of the behavioral studies, mice were sacrificed and counts of ventral MCs performed.

**Figure 1.**
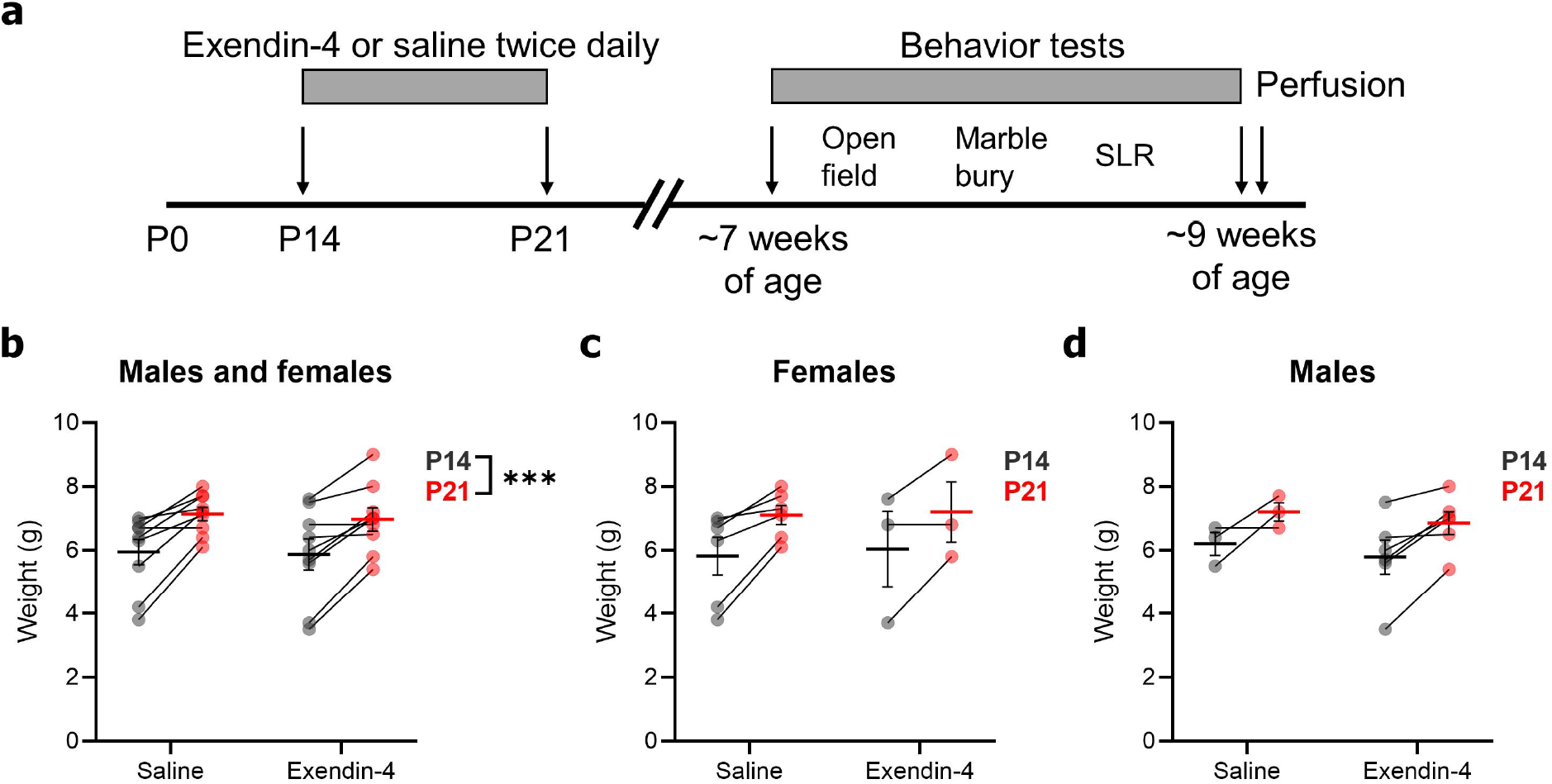
Experimental timeline. **(a)** Mice were treated with exendin-4 (0.5 mg/kg, dissolved in saline) or saline twice daily intraperitoneally from P14 through P21. Mice were subsequently weaned and allowed to develop uninterrupted. Starting at ∼7 weeks of age, mice were tested in open field, marble burying, and spontaneous location recognition (SLR) tests. Upon completion, mice were perfused for immunohistochemistry. **(b)** High-dose exendin-4 treatment did not significantly alter weight gain compared to saline treatment from P14 to P21 (Saline: N = 6F/3M, exendin-4: N = 3F/6M. two-way mixed ANOVA: main effect of age: F(1,16) = 39.26, p < 0.001; main effect of treatment: F(1,16) = 0.059, p = 0.81; age x treatment interaction: F(1,16) = 0.059, p = 0.81). **(c)** Weight gain, females only. **(d)** Weight gain, males only.

### Behavioral effects in adult mice

In the open field test, we found that adult mice treated with exendin-4 during development traveled significantly less total distance than control mice (**Fig. 2a-c**). However, the amount of time spent in the center of the open field did not significantly differ (**Fig. 2d-f**). Consistent with reduced locomotor activity, exendin-4-treated mice buried significantly fewer marbles than control mice (**Fig. 2g-h**).

**Figure 2.**
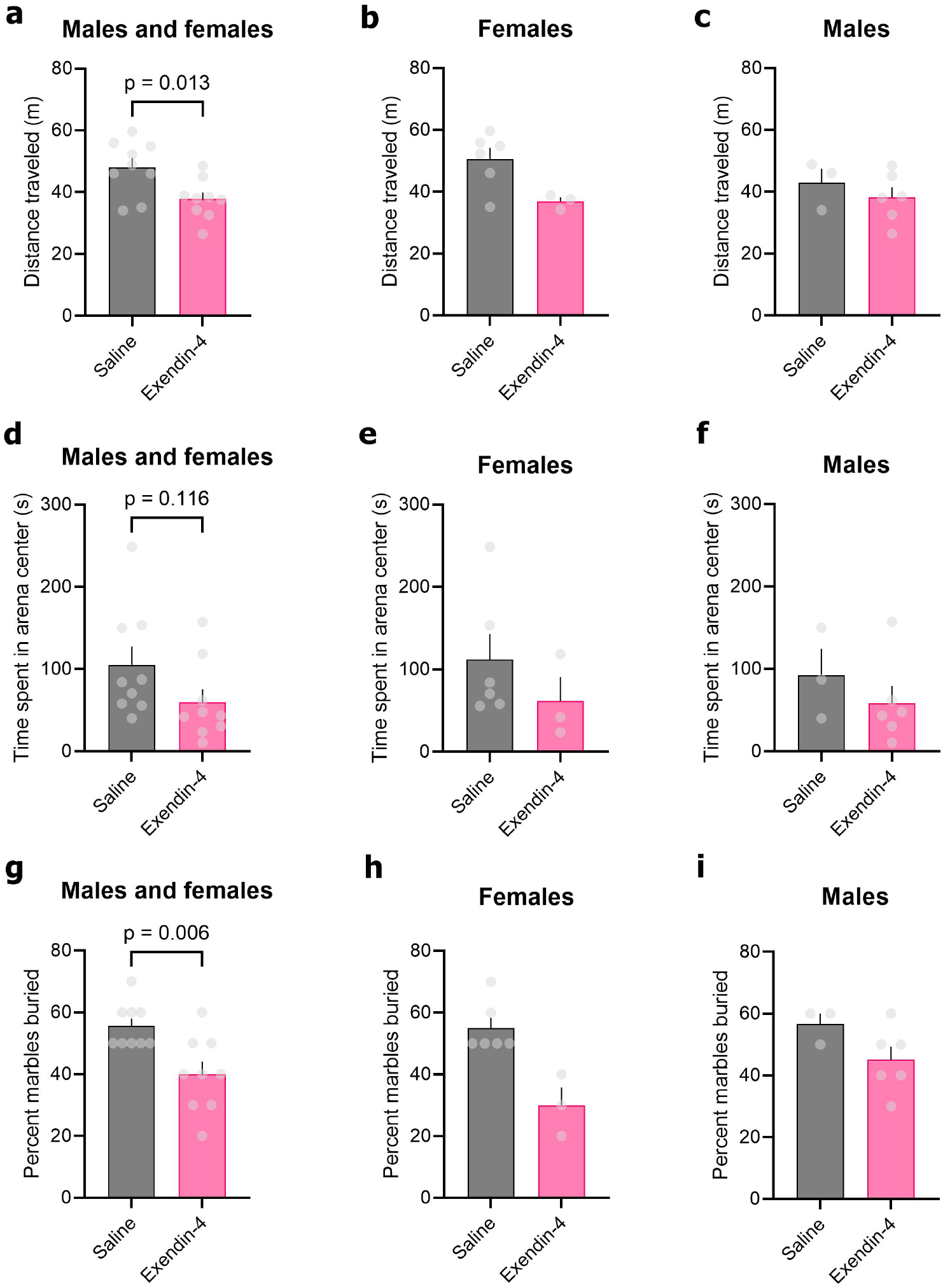
Effects of developmental exendin-4 treatment on behavioral tests of motor activity in adulthood. **(a)** Exendin-4 treatment reduces adult distance traveled in the open field test (unpaired t test: t(14.65) = 2.81, p = 0.013). **(b)** Distance traveled, females only. **(c)** Distance traveled, males only. **(d)** Time spent in the center of the open field did not significantly differ between exendin-4 and saline treated mice (unpaired t test: t(14.48) = 1.67, p = 0.12). **(e)** Center time, females only. **(f)** Center time, males only. **(g)** Exendin-4 treatment reduces percentage of marbles buried (unpaired t test: t(13.01) = 3.28, p = 0.006). **(h)** Marbles buried, females only. **(i)** Marbles buried, males only. For all experiments: saline: N = 6F/3M, exendin-4: N = 3F/6M.

In the SLR test (**Fig. 3a**), we did not detect significant differences in DI between control and exendin-4 treated mice (**Fig. 3b-d**). Importantly, both groups showed DIs significantly greater the 0, confirming intact pattern separation and memory. Both groups also spent similar times examining the objects in the testing arena (**Fig. 3e-g**), suggesting no differences in investigative motivation. Finally, we performed the SLR test using a mouse model of the 22q11.2 deletion syndrome, which has previously been shown to have impairments in hippocampal-dependent spatial learning and memory [23] and as such serves as a positive control for our task.

**Figure 3.**
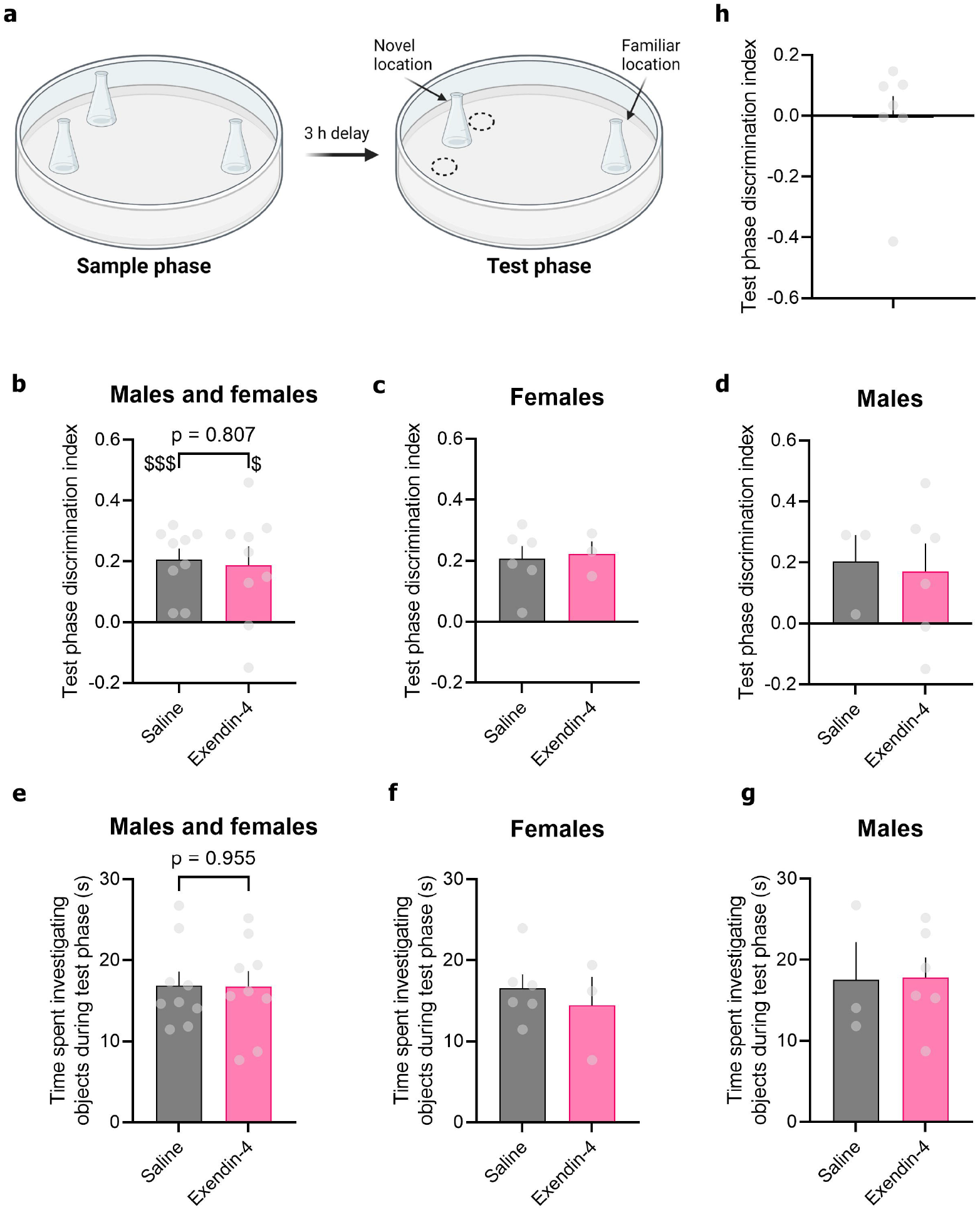
Effects of developmental exendin-4 treatment on hippocampal-dependent learning and memory function. **(a)** Schematic of the spontaneous location recognition (SLR) task. During the sample phase, three identical objects are placed within a circular arena, and mice allowed 10 mins to explore these objects. They are then returned to their home cage, and 3 h later placed back in the arena for the test phase. In the 5 min test phase, one object has remained in its original familiar location. The other original two objects have been removed (their original locations are marked in the schematic by dotted circles), and an identical object is placed in between the original locations (novel location). Because mice spontaneously prefer novelty, their ability to discriminate and commit to memory the object locations presented during the sample phase is reflected in a greater time spent exploring the object in the novel location than in the familiar location. **(b)** Mice treated with saline (N = 6F/3M) or exendin-4 (N = 3F/6M) both show significant discrimination of the novel location from the familiar location (one-sample t test versus discrimination index (DI) of 0: saline: t(8) = 5.58, p < 0.001, exendin-4: t(8) = 3.08, p = 0.015), but DI did not significantly differ between the treatment groups (unpaired t test: t(13.15) = 0.25, p = 0.81). $p < 0.05, $$$p < 0.001 in one-sample t test versus DI = 0. **(c)** Discrimination index, females only. **(d)** Discrimination index, males only. **(e)** Time spent investigating the objects during the test phase did not differ between saline (N = 6F/3M) or exendin-4 (N = 3F/6M) treatment groups (unpaired t test: t(15.79) = 0.057, p = 0.96). **(f)** Object investigation, females only. **(g)** Object investigation, males only. **(h)** As a positive control for the SLR test to identify hippocampal-dependent learning and memory deficits, we tested 22q11.2 deletion syndrome model mice (N = 7) of matched age to the exendin-4 and saline-treated mice. As expected, these mice were unable to discriminate between the novel and familiar locations (one-sample t test versus DI = 0: t(6) = 0.086, p = 0.93).

Discrimination index in 22q11.2 deletion syndrome mice did not differ from 0, suggesting that the SLR task is sensitive to hippocampal learning and memory deficits in our hands (**Fig. 3h**).

### Mossy cell effects in adult mice

In the mouse ventral hippocampus, the majority of *Glp1r* gene expression is detected in large glutamatergic neurons in the DG hilus call MCs, with only very minor *Glp1r* expression in a subset of neurons in area CA3 [17]. Ventral MCs from mice during early postnatal development show increased firing rate and resting membrane depolarization in response to bath application of exendin-4 [17]. Because of this functional expression of GLP-1R in ventral MCs, we hypothesized this population may be sensitive to deleterious effects of high-dose GLP-1R agonist during development. We used two different markers to detected MCs amongst neurons in the ventral DG hilus of adult mice: GluR2/3, which is expressed in hilar MCs across the hippocampal long axis [24, 25], and calretinin, which is specific for hilar MCs only in the ventral DG [26, 27]. As expected, GluR2/3 and calretinin overlapped extensively in ventral sections (**Fig. 4a**). However, we did not detect any significant differences in MC density between mice treated with exendin-4 or saline during development for marker (**Fig. 4b-g**).

**Figure 4.**
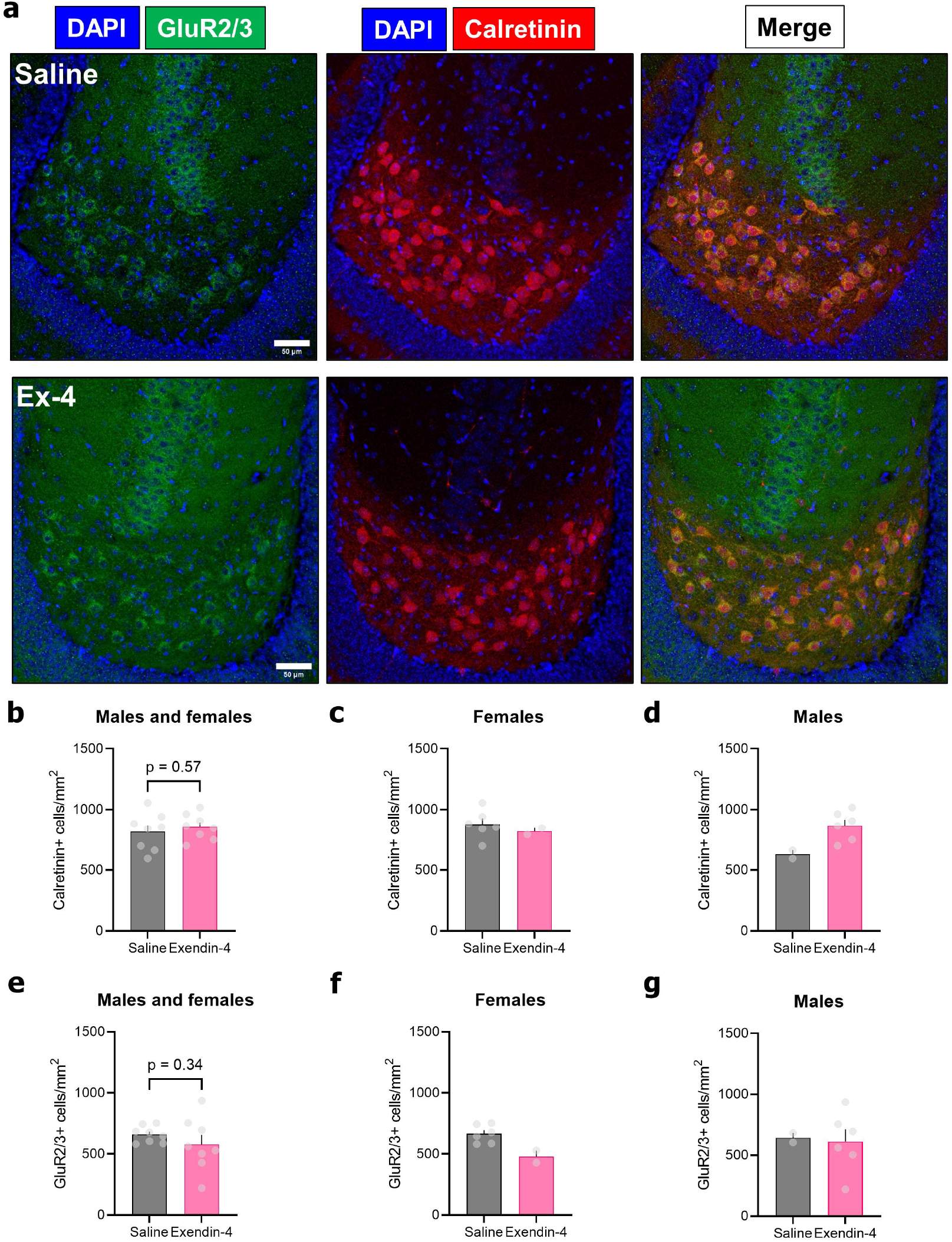
Effects of developmental exendin-4 treatment on hippocampal mossy cell (MC) number. **(a)** Mossy cells in the ventral dentate gyrus hilus were marked by expression of GluR2/3 (green) or calretinin (red) followed by confocal microscopy, demonstrating extensive overlap. Scale bar = 50 microns. **(b, e)** Developmental exendin-4 treatment did not significantly alter the number of ventral MCs as marked by calretinin ((b), unpaired t test: t(12.48) = 0.59, p = 0.57) or GluR2/3 ((e), unpaired t test: t(8.33) = 1.00, p = 0.34). **(c)** Calretinin-positive MCs, females only. **(d)** Calretinin-positive MCs, males only. **(f)** GluR2/3-positive MCs, females only. **(g)** GluR2/3-positive MCs, males only. For all experiments: saline: N = 6F/2M, exendin-4: N = 2F/6M.

## DISCUSSION

The main findings of this study were that high-dose exendin-4 administered twice daily beginning in the third postnatal week and continued through P21 1) does not alter weight gain during the drug administration period, 2) modestly reduces behavioral measures of motor activity, including total distance traveled in the open field test and percentage of marbles buried in adult mice, 3) does not significantly change adult performance in the SLR task, a sensitive measure of hippocampal-dependent pattern separation and learning and memory, and 4) does not change the number of adult ventral hippocampal MCs, which strongly express GLP-1R in mice. Finally, while this study was not powered to formally detect sex differences, disaggregation of the data between females and males did not suggest substantial effects of sex on any of the outcomes.

Understanding the developmental effects of a drug is challenging. The widespread expression of GLP-1R in the brain [28], coupled with the existence of critical or sensitive periods necessary for normal brain development, yields an almost infinite number of permutations between dosage, dose timing during development, age at subsequent testing, and selection of behavioral or structural outcomes. There is nothing to say that no behavioral effect observed with one dosing paradigm definitively means no behavioral effect would be observed for another.

However, by selecting a GLP-1R agonist, exendin-4, that has acute and/or chronic effects on adult mouse hippocampal activity, neuroprotection, adult neurogenesis, and cognition [17, 29-31] and administering it at a high dose during a period of massive brain development, our goal was to establish a focused experimental setup with high sensitivity for detecting effects that might narrow the scope of future, increasingly mechanistically oriented studies.

The most important of our findings is that despite very high-dose exposure to exendin-4 during development, behavioral effects in adulthood were focused rather than global and modest in size, consistent with a previous study of in utero exposure to exendin-4 in mice [11]. For instance, while exendin-4-treated mice traveled less distance in the open field, this is unlikely to be reflective of global listlessness, as they demonstrated comparable amounts of time investigating objects in the SLR test. Additionally, fewer marbles were buried by exendin-4-treated mice, though we fully agree with previous literature urging not to ascribe specific meaning to marble burying aside from a measure of activity which complements that of other assays like the open field test [32]. Considering the absence of investigative and cognitive effects in the SLR, it cannot be concluded as to whether the motor effects observed should be considered “pathological” at all. What they do suggest is that developmental effects of GLP-1R activation may be nuanced, influencing motivational and reward processes that underlie locomotor behavior, entirely consistent with the recognized role for GLP-1Rs and agonists, including exendin-4, on these processes in adult rodents [33-36].

Regarding structural changes, we focused our quantification on MCs of the ventral hippocampus because we felt this population might again provide a high likelihood of detecting changes stemming from developmental exposure to GLP-1R agonist. We recently demonstrated that ventral MCs are activated by acute systemic injection of exendin-4 at a dose much lower than used in the present study and that acute slices generated from pups in their first postnatal month of life respond to bath application of exendin-4 by increasing their firing rate and depolarizing their resting membrane potential [17]. Mossy cells, especially ventral MCs, are vulnerable to excitotoxic cell death after many types of brain insults or injuries (reviewed in [37]). As such, we hypothesized that repeated exposure to exendin-4 during development might chronically elevate ventral MC activity and contribute to developmental cell death. Our results did not suggest this was the case. This finding is interesting to compare with another study of early postnatal exendin-4 exposure (P0-P6) which found changes in hypothalamic circuitry [16]. Despite no loss of ventral MCs, MC function may still be altered. However, given the reliance of spatial pattern separation and memory on MC function [19, 38], results from the SLR test argue against this possibility as well. One other intriguing notion is that excitotoxic mechanisms from chronic ventral MC activation could be balanced by hippocampal neuroprotective mechanisms from GLP-1R activation, such as enhanced neurotrophin release (reviewed in [39]).

There are several limitations that must be considered when interpreting our results and in planning for future directions. First, as noted above, the study employed only one dosing paradigm for exendin-4, chosen for its previously demonstrated tolerability in early postnatal mice and its therapeutic effects on neonatal brain injury [12]. Future work must incorporate different GLP-1R agonists with dose ranges and additional dosing windows during development. Second, the generalizability of these findings across species must be considered as there are distinct differences in GLP-1R expression between species, for instance, expression in rat ventral CA1 neurons [40, 41] while *Glp1r* mRNA is not readily detected in mouse ventral CA1 pyramidal neurons [17, 42]. Third, our study only focused on exogenous GLP-1R agonists, whereas it remains unknown how endogenous GLP-1 release within the brain shapes brain development. Finally, additional investigation of sex differences will be important, as our study had relatively small and sample sizes of each sex and thus was not able to provide definitive conclusions on their presence or absence.

## CONCLUSION

In this study, high-dose exendin-4 administered from P14 to P21 significantly reduced open field locomotor activity and marble burying during adulthood as compared to saline treated mice, without changing performance on a behavioral task reflective of hippocampal-dependent learning and memory or changing the number of GLP-1R expressing MCs in the hippocampus. These findings support future investigation of how early GLP-1R activation influences (or does not influence) structure and function of brain regions important for cognition and motivated behavior.

## Acknowledgements

Figure 3a was created with BioRender.com.

